## Supplementary Table 1 for "An Endocytic Capture Model for Skeletal Muscle T-tubule Formation"

Supplementary Table 1. DNA constructs used in this study and Addgene identifiers

| **Vector name** | **Reference** | **Repository Identifier** |
|---|---|---|
| pME-2PH-PLCd | 46 | Addgene 71283 |
| pME-actn2 | This study | Addgene 109550 |
| pME-BTK-PH | 46 | Addgene 67673 |
| pME-Cav3 | This study | Addgene 109570 |
| pME-Cavin4a | This study | Addgene 109653 |
| pME-Cavin4b | This study | Addgene 109652 |
| pME-CD44a | This study | Addgene 109575 |
| pME-CD44b | This study | Addgene 109576 |
| pME-EGFP-NS | 41 | Addgene 75341 |
| pME-GalT | This study | Addgene 109558 |
| pME-lamp1 | This study | Addgene 109559 |
| pME-Lifeact | This study | Addgene 109545 |
| pME-mKate2-GPI-pA | This study | Addgene 109542 |
| pME-mKate2-KDEL-pA | This study | Addgene 109546 |
| pME-mKate2-NS | This study | Addgene 109729 |
| pME-mScarlet-NS | This study | Addgene 140874 |
| pME-mScarlet-STOP | This study | Addgene 140876 |
| pME-myristo | This study | Addgene 109541 |
| pME-nls-transposase-nls | This study | Addgene 140869 |
| pME-TCAP | This study | Addgene 109549 |
| p3E-2xFYVE | 46 | Addgene 109557 |
| p3E-ARF1 | This study | Addgene 109589 |
| p3E-ARF1-DN | This study | Addgene 109590 |
| p3E-ARF6 | This study | Addgene 109591 |
| p3E-ARF6-DN | This study | Addgene 109592 |
| p3E-ATG18 | This study | Addgene 109556 |
| p3E-Bin1a | This study | Addgene 109588 |
| p3E-Bin1b | This study | Addgene 109561 |
| p3E-CaaX(tH) | This study | Addgene 109539 |
| p3E-CaaX(tK) | This study | Addgene 109540 |
| p3E-CDC42 | This study | Addgene 109583 |
| p3E-CDC42-DN | This study | Addgene 109584 |
| p3E-Clta | This study | Addgene 109585 |
| p3E-Cltb | This study | Addgene 109586 |
| p3E-Dnm2a | This study | Addgene 109571 |
| p3E-Dnm2a-DN | This study | Addgene 109572 |
| p3E-Dnm2b | This study | Addgene 109573 |
| p3E-Dnm2b-DN | This study | Addgene 109574 |
| p3E-DysF | This study | Addgene 109544 |
| p3E-EEA1 | This study | Addgene 109560 |

| | | |
|---|---|---|
| p3E-EGFP | This study | Addgene 140877 |
| p3E-EHD1a | This study | Addgene 109564 |
| p3E-EHD1a-dEH | This study | Addgene 109565 |
| p3E-EHD1b | This study | Addgene 109566 |
| p3E-EHD1b-dEH | This study | Addgene 109567 |
| p3E-EHD2a | This study | Addgene 109568 |
| p3E-EHD2b | This study | Addgene 109569 |
| p3E-FAPP1 | This study | Addgene 109555 |
| p3E-ING2 | This study | Addgene 109554 |
| p3E-jph1a | This study | Addgene 109551 |
| p3E-Lact-C2 | This study | Addgene 109553 |
| p3E-mKate2 | This study | Addgene 109730 |
| p3E-mScarlet | This study | Addgene 140875 |
| p3E-MTM1 | This study | Addgene 109543 |
| p3E-pA | 40 | N/A |
| p3E-Pacsin2 | This study | Addgene 109587 |
| p3E-SERCA1 | This study | Addgene 109552 |
| p3E-rab1aa | This study | Addgene 109673 |
| p3E-rab1ab | This study | Addgene 109674 |
| p3E-rab1ba | This study | Addgene 109675 |
| p3E-rab1bb | This study | Addgene 109676 |
| p3E-rab2a | This study | Addgene 109687 |
| p3E-rab3aa | This study | Addgene 109703 |
| p3E-rab3ab | This study | Addgene 109704 |
| p3E-Rab3b | This study | Addgene 109752 |
| p3E-rab3c | This study | Addgene 109705 |
| p3E-rab3da | This study | Addgene 109706 |
| p3E-rab3db | This study | Addgene 109707 |
| p3E-rab4a | This study | Addgene 109714 |
| p3E-rab4a-DN | This study | Addgene 109735 |
| p3E-rab4a-CA | This study | Addgene 109736 |
| p3E-rab4b | This study | Addgene 109715 |
| p3E-rab5aa | This study | Addgene 109716 |
| p3E-rab5ab | This study | Addgene 109717 |
| p3E-rab5b | This study | Addgene 109718 |
| p3E-rab5c | This study | Addgene 109719 |
| p3E-rab5c-DN | This study | Addgene 109737 |
| p3E-rab5c-CA | This study | Addgene 109738 |
| p3E-rab6a | This study | Addgene 109720 |
| p3E-rab6a-DN | This study | Addgene 145303 |
| p3E-rab6a-CA | This study | Addgene 109733 |
| p3E-rab6ba | This study | Addgene 109721 |
| p3E-rab6bb | This study | Addgene 109722 |

| | | |
|---|---|---|
| p3E-rab7a | This study | Addgene 109723 |
| p3E-rab7b | This study | Addgene 109724 |
| p3E-rab8a | This study | Addgene 109725 |
| p3E-rab8a-DN | This study | Addgene 145305 |
| p3E-rab8a-CA | This study | Addgene 145304 |
| p3E-rab8b | This study | Addgene 109726 |
| p3E-rab9a | This study | Addgene 109727 |
| p3E-rab9b | This study | Addgene 109728 |
| p3E-rab10 | This study | Addgene 109661 |
| p3E-rab11a | This study | Addgene 109662 |
| p3E-rab11a-DN | This study | Addgene 109739 |
| p3E-rab11a-CA | This study | Addgene 109740 |
| p3E-rab11al | This study | Addgene 109663 |
| p3E-rab11ba | This study | Addgene 109664 |
| p3E-rab11ba-DN | This study | Addgene 145291 |
| p3E-rab11ba-CA | This study | Addgene 145290 |
| p3E-rab11bb | This study | Addgene 109665 |
| p3E-rab12 | This study | Addgene 109666 |
| p3E-rab13 | This study | Addgene 109667 |
| p3E-rab13-DN | This study | Addgene 145293 |
| p3E-rab13-CA | This study | Addgene 145292 |
| p3E-rab14 | This study | Addgene 109668 |
| p3E-rab14l | This study | Addgene 109669 |
| p3E-rab15 | This study | Addgene 109670 |
| p3E-rab18a | This study | Addgene 109671 |
| p3E-rab18b | This study | Addgene 109672 |
| p3E-Rab19 | This study | Addgene 109753 |
| p3E-rab20 | This study | Addgene 109677 |
| p3E-rab21 | This study | Addgene 109678 |
| p3E-rab22a | This study | Addgene 109679 |
| p3E-rab22a-DN | This study | Addgene 145295 |
| p3E-rab22a-CA | This study | Addgene 145294 |
| p3E-rab23 | This study | Addgene 109680 |
| p3E-rab23-DN | This study | Addgene 109734 |
| p3E-rab23-CA | This study | Addgene 145296 |
| p3E-rab24 | This study | Addgene 109681 |
| p3E-rab25a | This study | Addgene 109682 |
| p3E-rab25b | This study | Addgene 109683 |
| p3E-rab26 | This study | Addgene 109684 |
| p3E-rab26-DN | This study | Addgene 109745 |
| p3E-rab26-CA | This study | Addgene 109746 |
| p3E-rab27a | This study | Addgene 109685 |
| p3E-Rab27b | This study | Addgene 109754 |

| | | |
|---|---|---|
| p3E-rab28 | This study | Addgene 109686 |
| p3E-rab28-DN | This study | Addgene 145298 |
| p3E-rab28-CA | This study | Addgene 145927 |
| p3E-rab30 | This study | Addgene 109688 |
| p3E-Rab31 | This study | Addgene 109755 |
| p3E-rab32a | This study | Addgene 109689 |
| p3E-rab32b | This study | Addgene 109690 |
| p3E-rab32b-DN | This study | Addgene 145300 |
| p3E-rab32b-CA | This study | Addgene 145299 |
| p3E-rab33a | This study | Addgene 109691 |
| p3E-rab33ba | This study | Addgene 109692 |
| p3E-Rab33bb | This study | Addgene 109756 |
| p3E-rab33bb-DN | This study | Addgene 109741 |
| p3E-rab33bb-CA | This study | Addgene 109742 |
| p3E-rab34a | This study | Addgene 109693 |
| p3E-rab34b | This study | Addgene 109694 |
| p3E-rab35a | This study | Addgene 109695 |
| p3E-rab35b | This study | Addgene 109696 |
| p3E-rab36 | This study | Addgene 109697 |
| p3E-rab37 | This study | Addgene 109698 |
| p3E-rab37-DN | This study | Addgene 109743 |
| p3E-rab37-CA | This study | Addgene 109744 |
| p3E-rab38a | This study | Addgene 109699 |
| p3E-rab38a-DN | This study | Addgene 109747 |
| p3E-rab38a-CA | This study | Addgene 109748 |
| p3E-Rab38b | This study | Addgene 109757 |
| p3E-Rab38c | This study | Addgene 109758 |
| p3E-rab39a | This study | Addgene 109700 |
| p3E-rab39ba | This study | Addgene 109701 |
| p3E-rab39bb | This study | Addgene 109702 |
| p3E-rab40b | This study | Addgene 109708 |
| p3E-rab40b-DN | This study | Addgene 145302 |
| p3E-rab40b-CA | This study | Addgene 145301 |
| p3E-rab40c | This study | Addgene 109709 |
| p3E-Rab41 | This study | Addgene 109759 |
| p3E-rab42a | This study | Addgene 109710 |
| p3E-rab42b | This study | Addgene 109711 |
| p3E-rab43 | This study | Addgene 109712 |
| p3E-rab44 | This study | Addgene 109713 |
| actb2-lifeact-mKate2 | This study | Addgene 109750 |
| actb2-mKate2-KDEL-pA | This study | Addgene 109749 |
| actc1b- mKate2-KDEL-pA | This study | Addgene 109494 |
| actc1b-2PH-PLCd-mKate2 | This study | Addgene 109500 |

| | | |
|---|---|---|
| actc1b-actn2-mKate2 | This study | Addgene 109751 |
| actc1b-BTK-mKate2 | This study | Addgene 109501 |
| actc1b-Cav3-mKate2 | This study | Addgene 109518 |
| actc1b-Cavin4a-mKate2 | This study | Addgene 109510 |
| actc1b-Cavin4b-mKate2 | This study | Addgene 109511 |
| actc1b-CD44a-mKate2 | This study | Addgene 109523 |
| actc1b-CD44b-mKate2 | This study | Addgene 109524 |
| actc1b-GalT-mKate2 | This study | Addgene 109506 |
| actc1b-lamp1-mKate2 | This study | Addgene 109507 |
| actc1b-Lifeact-mKate2 | This study | Addgene 109493 |
| actc1b-mKate2-2xFYVEhgs | This study | Addgene 109505 |
| actc1b-mKate2-ARF1 | This study | Addgene 109535 |
| actc1b-mKate2-ARF1-DN | This study | Addgene 109536 |
| actc1b-mKate2-ARF6 | This study | Addgene 109537 |
| actc1b-mKate2-ARF6-DN | This study | Addgene 109538 |
| actc1b-mKate2-ATG18 | This study | Addgene 109504 |
| actc1b-mKate2-Bin1a | This study | Addgene 109534 |
| actc1b-mKate2-Bin1b | This study | Addgene 109509 |
| actc1b-mKate2-CaaX(tH) | This study | Addgene 109487 |
| actc1b-mKate2-CaaX(tK) | This study | Addgene 109488 |
| actc1b-mKate2-CDC42 | This study | Addgene 109531 |
| actc1b-mKate2-CDC42-DN | This study | Addgene 109532 |
| actc1b-mKate2-Dnm2a | This study | Addgene 109519 |
| actc1b-mKate2-Dnm2a-DN | This study | Addgene 109520 |
| actc1b-mKate2-Dnm2b | This study | Addgene 109521 |
| actc1b-mKate2-Dnm2b-DN | This study | Addgene 109522 |
| actc1b-mKate2-DysF | This study | Addgene 109492 |
| actc1b-mKate2-EEA1 | This study | Addgene 109508 |
| actc1b-mKate2-EHD1a | This study | Addgene 109512 |
| actc1b-mKate2-EHD1a-dEH | This study | Addgene 109513 |
| actc1b-mKate2-EHD1b | This study | Addgene 109514 |
| actc1b-mKate2-EHD1b-dEH | This study | Addgene 109515 |
| actc1b-mKate2-EHD2a | This study | Addgene 109516 |
| actc1b-mKate2-EHD2b | This study | Addgene 109517 |
| actc1b-mKate2-FAPP1 | This study | Addgene 109503 |
| actc1b-mKate2-GPI-pA | This study | Addgene 109490 |
| actc1b-mKate2-ING2 | This study | Addgene 109502 |
| actc1b-mKate2-jph1a | This study | Addgene 109497 |
| actc1b-mKate2-LactC2 | This study | Addgene 109499 |
| actc1b-mKate2-MTM1 | This study | Addgene 109491 |
| actc1b-mKate2-Pacsin2 | This study | Addgene 109533 |
| actc1b-mKate2-SERCA1 | This study | Addgene 109498 |
| actc1b-myristo-mKate2 | This study | Addgene 109489 |

| | | |
|---|---|---|
| actc1b-TCAP-mKate2 | This study | Addgene 109496 |
| actc1b-mKate2-rab1aa | This study | Addgene 109605 |
| actc1b-mKate2-rab1ab | This study | Addgene 109606 |
| actc1b-mKate2-rab1ba | This study | Addgene 109607 |
| actc1b-mKate2-rab1bb | This study | Addgene 109608 |
| actc1b-mKate2-rab2a | This study | Addgene 109619 |
| actc1b-mKate2-rab3aa | This study | Addgene 109635 |
| actc1b-mKate2-rab3ab | This study | Addgene 109636 |
| actc1b-mKate2-Rab3b | This study | Addgene 109760 |
| actc1b-mKate2-rab3c | This study | Addgene 109637 |
| actc1b-mKate2-rab3da | This study | Addgene 109638 |
| actc1b-mKate2-rab3db | This study | Addgene 109639 |
| actc1b-mKate2-rab4a | This study | Addgene 109646 |
| actc1b-mKate2-rab4a-DN | This study | Addgene 145315 |
| actc1b-mKate2-rab4a-CA | This study | Addgene 145316 |
| actc1b-mKate2-rab4b | This study | Addgene 109647 |
| actc1b-mKate2-rab5aa | This study | Addgene 109648 |
| actc1b-mKate2-rab5ab | This study | Addgene 109649 |
| actc1b-mKate2-rab5b | This study | Addgene 109650 |
| actc1b-mKate2-rab5c | This study | Addgene 109651 |
| actc1b-mKate2-rab5c-DN | This study | Addgene 145319 |
| actc1b-mKate2-rab5c-CA | This study | Addgene 145320 |
| actc1b-mKate2-rab6a | This study | Addgene 109652 |
| actc1b-mKate2-rab6a-DN | This study | Addgene 145307 |
| actc1b-mKate2-rab6a-CA | This study | Addgene 145308 |
| actc1b-mKate2-rab6ba | This study | Addgene 109653 |
| actc1b-mKate2-rab6bb | This study | Addgene 109654 |
| actc1b-mKate2-rab7a | This study | Addgene 109655 |
| actc1b-mKate2-rab7b | This study | Addgene 109656 |
| actc1b-mKate2-rab8a | This study | Addgene 109657 |
| actc1b-mKate2-rab8a-DN | This study | Addgene 145337 |
| actc1b-mKate2-rab8a-CA | This study | Addgene 145338 |
| actc1b-mKate2-rab8b | This study | Addgene 109658 |
| actc1b-mKate2-rab8b-DN | This study | Addgene 145371 |
| actc1b-mKate2-rab8b-CA | This study | Addgene 145372 |
| actc1b-mKate2-rab9a | This study | Addgene 109659 |
| actc1b-mKate2-rab9b | This study | Addgene 109660 |
| actc1b-mKate2-rab10 | This study | Addgene 109593 |
| actc1b-mKate2-rab11a | This study | Addgene 109594 |
| actc1b-mKate2-rab11a-DN | This study | Addgene 145313 |
| actc1b-mKate2-rab11a-CA | This study | Addgene 145314 |
| actc1b-mKate2-rab11al | This study | Addgene 109595 |
| actc1b-mKate2-rab11ba | This study | Addgene 109596 |

| | | |
|---|---|---|
| actc1b-mKate2-rab11ba-DN | This study | Addgene 145311 |
| actc1b-mKate2-rab11ba-CA | This study | Addgene 145312 |
| actc1b-mKate2-rab11bb | This study | Addgene 109597 |
| actc1b-mKate2-rab12 | This study | Addgene 109598 |
| actc1b-mKate2-rab13 | This study | Addgene 109599 |
| actc1b-mKate2-rab13-DN | This study | Addgene 145329 |
| actc1b-mKate2-rab13-CA | This study | Addgene 145330 |
| actc1b-mKate2-rab14 | This study | Addgene 109600 |
| actc1b-mKate2-rab14l | This study | Addgene 109601 |
| actc1b-mKate2-rab15 | This study | Addgene 109602 |
| actc1b-mKate2-rab18a | This study | Addgene 109603 |
| actc1b-mKate2-rab18b | This study | Addgene 109604 |
| actc1b-mKate2-Rab19 | This study | Addgene 109761 |
| actc1b-mKate2-rab20 | This study | Addgene 109609 |
| actc1b-mKate2-rab21 | This study | Addgene 109610 |
| actc1b-mKate2-rab22a | This study | Addgene 109611 |
| actc1b-mKate2-rab22a-DN | This study | Addgene 145317 |
| actc1b-mKate2-rab22a-CA | This study | Addgene 145318 |
| actc1b-mKate2-rab23 | This study | Addgene 109612 |
| actc1b-mKate2-rab23-DN | This study | Addgene 145309 |
| actc1b-mKate2-rab23-CA | This study | Addgene 145310 |
| actc1b-mKate2-rab24 | This study | Addgene 109613 |
| actc1b-mKate2-rab25a | This study | Addgene 109614 |
| actc1b-mKate2-rab25b | This study | Addgene 109615 |
| actc1b-mKate2-rab26 | This study | Addgene 109616 |
| actc1b-mKate2-rab26-DN | This study | Addgene 145333 |
| actc1b-mKate2-rab26-CA | This study | Addgene 145334 |
| actc1b-mKate2-rab27a | This study | Addgene 109617 |
| actc1b-mKate2-Rab27b | This study | Addgene 109762 |
| actc1b-mKate2-rab28 | This study | Addgene 109618 |
| actc1b-mKate2-rab28-DN | This study | Addgene 145321 |
| actc1b-mKate2-rab28-CA | This study | Addgene 145322 |
| actc1b-mKate2-rab30 | This study | Addgene 109620 |
| actc1b-mKate2-rab31 | This study | Addgene 109763 |
| actc1b-mKate2-rab32a | This study | Addgene 109621 |
| actc1b-mKate2-rab32b | This study | Addgene 109622 |
| actc1b-mKate2-rab32b-DN | This study | Addgene 145325 |
| actc1b-mKate2-rab32b-CA | This study | Addgene 145326 |
| actc1b-mKate2-rab33a | This study | Addgene 109623 |
| actc1b-mKate2-rab33ba | This study | Addgene 109624 |
| actc1b-mKate2-rab33bb | This study | Addgene 109764 |
| actc1b-mKate2-rab33bb-DN | This study | Addgene 145323 |
| actc1b-mKate2-rab33bb-CA | This study | Addgene 145324 |

| | | |
|---|---|---|
| actc1b-mKate2-rab34a | This study | Addgene 109625 |
| actc1b-mKate2-rab34b | This study | Addgene 109626 |
| actc1b-mKate2-rab35a | This study | Addgene 109627 |
| actc1b-mKate2-rab35b | This study | Addgene 109628 |
| actc1b-mKate2-rab36 | This study | Addgene 109629 |
| actc1b-mKate2-rab37 | This study | Addgene 109630 |
| actc1b-mKate2-rab37-DN | This study | Addgene 145331 |
| actc1b-mKate2-rab37-CA | This study | Addgene 145332 |
| actc1b-mKate2-rab38a | This study | Addgene 109631 |
| actc1b-mKate2-rab38a-DN | This study | Addgene 145335 |
| actc1b-mKate2-rab38a-CA | This study | Addgene 145336 |
| actc1b-mKate2-Rab38b | This study | Addgene 109765 |
| actc1b-mKate2-Rab38c | This study | Addgene 109766 |
| actc1b-mKate2-rab39a | This study | Addgene 109632 |
| actc1b-mKate2-rab39ba | This study | Addgene 109633 |
| actc1b-mKate2-rab39bb | This study | Addgene 109634 |
| actc1b-mKate2-rab40b | This study | Addgene 109640 |
| actc1b-mKate2-rab40b-DN | This study | Addgene 145327 |
| actc1b-mKate2-rab40b-CA | This study | Addgene 145328 |
| actc1b-mKate2-rab40c | This study | Addgene 109641 |
| actc1b-mKate2-Rab41 | This study | Addgene 109767 |
| actc1b-mKate2-rab42a | This study | Addgene 109642 |
| actc1b-mKate2-rab42b | This study | Addgene 109643 |
| actc1b-mKate2-rab43 | This study | Addgene 109644 |
| actc1b-mKate2-rab44 | This study | Addgene 109645 |
| mScarlet-bin1b in pCSDEST2 | This study | Addgene 145306 |
| Cav3-EGFP in pCSDEST2 | This study | Addgene 140889 |
| EGFP-CaaX(tH) in pCSDEST2 | This study | Addgene 140890 |
| EGFP-Rab8a in pCSDEST2 | This study | Addgene 140880 |
| EGFP-Rab8b in pCSDEST2 | This study | Addgene 140881 |
| EGFP-Rab13 in pCSDEST2 | This study | Addgene 140879 |
| EGFP-Rab19 in pCSDEST2 | This study | Addgene 140888 |
| EGFP-Rab26 in pCSDEST2 | This study | Addgene 140884 |
| EGFP-Rab32b in pCSDEST2 | This study | Addgene 140885 |
| EGFP-Rab33bb in pCSDEST2 | This study | Addgene 140882 |
| EGFP-Rab37 in pCSDEST2 | This study | Addgene 140886 |
| EGFP-Rab38a in pCSDEST2 | This study | Addgene 140887 |
| EGFP-Rab40b in pCSDEST2 | This study | Addgene 140883 |
| EGFP-SNX8 (mouse) in pEGFP-C1 | 30 | N/A |
| pT3TS_Transposase | This study | Addgene 109768 |
| nls-transposase-nls-pA in pT3TS-Dest | This study | Addgene 140873 |
| pDONR_p1-p3 | This study | Addgene 140865 |
| pCSDEST2 | 39 | Addgene 22424 |

| | | |
|---|---|---|
| pDEST-actb2-ccdb | This study | Addgene 109770 |
| pDEST-actc1b-ccdb | This study | Addgene 109771 |
| pDEST-actc1b-ccdb-mKate2 | This study | Addgene 109731 |
| pDEST-actc1b-mKate2-ccdb | This study | Addgene 109732 |
| pDEST-Tol2-pA2 | 40 | N/A |
| pT3TS-Dest_R1-R3 | This study | Addgene 140878 |
| pT7gRNA-ccdb | This study | Addgene 140864 |
| pT7gRNA-smyhc1_977 | This study | Addgene 140867 |
| pT7gRNA-ttn.1_C3 | This study | Addgene 140866 |
| pT7tyrgRNA | 51 | Addgene 46761 |
