## Supplementary Macro Code for "An Endocytic Capture Model for Skeletal Muscle T-tubule Formation"

### Macro 1 “GEN_Split_to_tif_CZI_AND_LSM.ijm

```
//TAKES .LSM OR CZI FILES AND SPLITS INTO DIFFERENT FILES FOR THE SINGLE CHANNELS AND A
MERGE

// SPECIFYING DIRECTORY, LISTING INPUT FILES AND MAKING RESULTS FOLDER
Imagepath = getDirectory("Choose a Directory containing lsm/czi files");
list = getFileList(Imagepath);

//SETTING UP LOG FILE
 print("\\Clear");
getDateAndTime(year, month, dayOfWeek, dayOfMonth, hour, minute, second, msec);

if (dayOfMonth<10) {
dayOfMonth = "0" + toString(dayOfMonth);
}

month = month+1;
if (month<10) {
month = "0" + month;
}

if (hour<10) {
hour = "0" + hour;
}

if (minute<10) {
minute = "0" + minute;
}

    ts = toString(dayOfMonth) + "-" + month + "-" + year + " " + hour + ":" + minute;
    IJ.log("STARTED macro: Split_to_tif_CZI_AND_LSM_v4.ijm, " + ts);

//SPECIFYING FILE EXTENSIONS
ImageEnd = ".lsm";
ImageEnd1 = ".czi";

// DIALOGUE BOX FOR INPUT
                    subfolder = "Split";
                    Dialog.create("Specify Variables");
                              Dialog.addString("New subfolder:", subfolder, 30);
                              Dialog.addCheckbox("Enhance Contrast (0.35% saturated pixels)", false);
                              Dialog.addCheckbox("Convert to 8-bit RGB", false);
                              Dialog.addCheckbox("Save single channels", true);
                              Dialog.addCheckbox("Save merge", true);
                              Dialog.addCheckbox("Use Channel Shuffler", false);
                              Dialog.addCheckbox("Notify about unprocessed stacks/multichannel files",
false);
                              Dialog.show();
                    subfolder = Dialog.getString();
                    EnhCont = Dialog.getCheckbox();
                    eightbit = Dialog.getCheckbox();
                    single = Dialog.getCheckbox();
                    merge = Dialog.getCheckbox();
                    shuffler = Dialog.getCheckbox();
```

```
                    unprocessed = Dialog.getCheckbox();

     resultsDir = Imagepath+subfolder+"/";
     File.makeDirectory(resultsDir);

    IJ.log("Input folder: " + Imagepath);
    IJ.log("Subfolder: " + subfolder);
                    if(EnhCont==1){
                    EnhCont_txt = "YES";
                    } else {EnhCont_txt = "NO";}
    IJ.log("Enhance Contrast (0.35% saturated pixels): " + EnhCont_txt);
                    if(eightbit==1)  {
                    eightbit_txt = "YES";
                    } else {eightbit_txt = "NO";}
    IJ.log("Convert to 8-bit RGB: " + eightbit_txt);
                    if(single==1)    {
                    single_txt = "YES";
                    } else {single_txt = "NO";}
    IJ.log("Save single channels: " + single_txt);
                    if(merge==1)   {
                    merge_txt = "YES";
                    } else {merge_txt = "NO";}
    IJ.log("Save merge: " + merge_txt);
                    if(shuffler==1)  {
                    shuffler_txt = "YES";
                    } else {shuffler_txt = "NO";}
    IJ.log("Use Channel Shuffler: " + shuffler_txt);
                    if(unprocessed==1)      {
                    unprocessed_txt = "YES";
                    } else {unprocessed_txt = "NO";}
    IJ.log("Notify about unprocessed stacks/multichannel files: " + unprocessed_txt);

//LOOP

for (i=0; i<list.length; i++)                                       {

// FOR LSM IMAGE FILES

 if (endsWith(list[i],ImageEnd))                            {

  open(Imagepath+list[i]);
  orig = getTitle();
  IJ.log("File opened: " + orig);
  nodotname = replace(orig,ImageEnd, "");

//CLOSE SINGLE CHANNEL AND 4+ CHANNEL IMAGES WITHOUT SAVING
getDimensions(width, height, channels, slices, frames);
   if (frames>1 || slices>1)        {
//CHANNEL NUMBER RESET NEEDED IN ORDER TO NOT CONFUSE LATER WITH 1 CHANNEL IMAGES
THAT HAVE MULTIPLE SLICES/FRAMES
   channels = 0;
   if (unprocessed==true)          {
waitForUser( "Input problem","This image is a stack");
                                                                    }
   IJ.log("***File is a stack, not saved");
  close();
                                                                    }
```

```
  if (channels>=4)                                              {
  if (unprocessed==true)                  {
waitForUser( "Input problem","This image has more than 3 channels");
                                                                {
  IJ.log("***File has >4 channels, not saved");
 close();
                                                                                }
  if (channels==1 && frames==1 && slices==1) {
                                        run("Grays");
                // ENHANCE CONTRAST
                                if (EnhCont==true)      {
                                        run("Enhance Contrast", "saturated=0.35");
                                                                        }

                // CONVERT TO 8-BIT RGB
                                if (eightbit==true)     {
                                        run("RGB Color");
                                                                        }
saveAs("tiff", resultsDir+nodotname+"_single_channel.tif");
  IJ.log("File saved: " + nodotname+"_single_channel.tif");
close();
                                                                                }
  if (channels==2 || channels ==3)  {

                // MANUAL CHANNEL SHUFFLER
                                if (shuffler==true)     {
                                        run("Arrange Channels...");
                                                                        }
 run("Split Channels");
 ChannelNo = nImages();


                // AUTO CHANNEL SHUFFLER - DEFAULT FOR TWO CHANNELS IS GR, THREE IS BRG
if (shuffler==false)        {

  if (ChannelNo==2)               {
     selectWindow("C1-"+orig);
                                        run("Green");
     selectWindow("C2-"+orig);
                                        run("Red");
                                                    }

  if (ChannelNo==3)               {
     selectWindow("C1-"+orig);
                                        run("Blue");
     selectWindow("C2-"+orig);
                                        run("Red");
     selectWindow("C3-"+orig);
                                        run("Green");
                                                    }
                                              }


 selectWindow("C1-"+orig);
                // ENHANCE CONTRAST
                                if (EnhCont==true)      {
                                        run("Enhance Contrast", "saturated=0.35");
```

```
                                                            }

  selectWindow("C2-"+orig);
                  // ENHANCE CONTRAST
                            if (EnhCont==true)      {
                                run("Enhance Contrast", "saturated=0.35");
                                                            }

   if (ChannelNo==3)             {
       selectWindow("C3-"+orig);
                  // ENHANCE CONTRAST
                            if (EnhCont==true)      {
                                run("Enhance Contrast", "saturated=0.35");
                                                            }
                                          }

//CHANNEL MERGING. FIJI DEFAULTS ARE: c1=RED, c2=GREEN, c3=BLUE
if (ChannelNo==2)                   {
run("Merge Channels...", "c1=C1-"+orig+" c2=C2-"+orig+" create keep");
                                                    }

if (ChannelNo==3)                   {
run("Merge Channels...", "c3=C3-"+orig+" c2=C2-"+orig+" c1=C1-"+orig+" create keep");
                                                    }


  selectWindow("C1-"+orig);
                  // CONVERT TO 8-BIT RGB
                            if (eightbit==true)       {
                                run("RGB Color");
                                                            }
if (single==true)               {
  saveAs("tiff", resultsDir+nodotname+"_C1.tif");
   IJ.log("File saved: " +nodotname+"_C1.tif");
                                        }
  close();

  selectWindow("C2-"+orig);
                  // CONVERT TO 8-BIT RGB
                            if (eightbit==true)       {
                                run("RGB Color");
                                                            }
if (single==true)               {
  saveAs("tiff", resultsDir+nodotname+"_C2.tif");
   IJ.log("File saved: " +nodotname+"_C2.tif");
                                        }
  close();

if (ChannelNo==3)                         {
          selectWindow("C3-"+orig);
                  // CONVERT TO 8-BIT RGB
                            if (eightbit==true)             {
                                run("RGB Color");
                                                                 }
if (single==true)               {
  saveAs("tiff", resultsDir+nodotname+"_C3.tif");
   IJ.log("File saved: " +nodotname+"_C3.tif");
```

```
                                                    }
  close();
                                                                          }

  selectWindow(orig);

                    // CONVERT TO 8-BIT RGB
                                    if (eightbit==true)        {
                                            run("RGB Color");
                                                                                   }
if (merge==true)                 {
  saveAs("tiff", resultsDir+nodotname+"_merge.tif");
   IJ.log("File saved: " +nodotname+"_merge.tif");
                                                    }
run("Close All");

                                                                                           }
} // END OF LSM LOOP



 if (endsWith(list[i],ImageEnd1))                            {  // START OF CZI LOOP

run("Bio-Formats  Importer",  "open="  +Imagepath+list[i]+"  color_mode=Default  view=Hyperstack
stack_order=XYCZT");
  orig = getTitle();
  IJ.log("File opened: " + orig);
  nodotname = replace(orig,ImageEnd1, "");

//CLOSE SINGLE CHANNEL AND 4+ CHANNEL IMAGES WITHOUT SAVING
getDimensions(width, height, channels, slices, frames);
   if (frames>1 || slices>1)        {
//CHANNEL NUMBER RESET NEEDED IN ORDER TO NOT CONFUSE LATER WITH 1 CHANNEL IMAGES
THAT HAVE MULTIPLE SLICES/FRAMES
   channels = 0;
   if (unprocessed==true)          {
waitForUser( "Input problem","This image is a stack");
                                                                  }
   IJ.log("***File is a stack, not saved");
  close();
                                                                  }
   if (channels>=4)                                                       {
   if (unprocessed==true)                  {
waitForUser( "Input problem","This image has more than 3 channels");
                                                                          }
   IJ.log("***File has >4 channels, not saved");
  close();
                                                                                           }
   if (channels==1 && frames==1 && slices==1) {
                                            run("Grays");
                    // ENHANCE CONTRAST
                                    if (EnhCont==true)       {
                                            run("Enhance Contrast", "saturated=0.35");
                                                                                   }

                    // CONVERT TO 8-BIT RGB
                                    if (eightbit==true)        {
```

```
				run("RGB Color");
								}
saveAs("tiff", resultsDir+nodotname+"_single_channel.tif");
   IJ.log("File saved: " + nodotname+"_single_channel.tif");
close();
										}
   if (channels==2 || channels ==3)  {

			// MANUAL CHANNEL SHUFFLER
				if (shuffler==true)	{
					run("Arrange Channels...");
								}
  run("Split Channels");
  ChannelNo = nImages();


			// AUTO CHANNEL SHUFFLER - DEFAULT FOR TWO CHANNELS IS GR, THREE IS BRG
if (shuffler==false)	{

   if (ChannelNo==2)		{
	selectWindow("C1-"+orig);
					run("Green");
	selectWindow("C2-"+orig);
					run("Red");
						}

   if (ChannelNo==3)		{
	selectWindow("C1-"+orig);
					run("Blue");
	selectWindow("C2-"+orig);
					run("Red");
	selectWindow("C3-"+orig);
					run("Green");
						}
					}


  selectWindow("C1-"+orig);
			// ENHANCE CONTRAST
				if (EnhCont==true)	{
					run("Enhance Contrast", "saturated=0.35");
								}

  selectWindow("C2-"+orig);
			// ENHANCE CONTRAST
				if (EnhCont==true)	{
					run("Enhance Contrast", "saturated=0.35");
								}

   if (ChannelNo==3)		{
	selectWindow("C3-"+orig);
			// ENHANCE CONTRAST
				if (EnhCont==true)	{
					run("Enhance Contrast", "saturated=0.35");
								}
						}
```

```
//CHANNEL MERGING. FIJI DEFAULTS ARE: c1=RED, c2=GREEN, c3=BLUE
if (ChannelNo==2)                    {
run("Merge Channels...", "c1=C1-"+orig+" c2=C2-"+orig+" create keep");
                                                                }

if (ChannelNo==3)                    {
run("Merge Channels...", "c3=C3-"+orig+" c2=C2-"+orig+" c1=C1-"+orig+" create keep");
                                                                }


  selectWindow("C1-"+orig);
              // CONVERT TO 8-BIT RGB
                          if (eightbit==true)       {
                                run("RGB Color");
                                                                          }
if (single==true)             {
  saveAs("tiff", resultsDir+nodotname+"_C1.tif");
   IJ.log("File saved: " +nodotname+"_C1.tif");
                                          }
  close();

  selectWindow("C2-"+orig);
              // CONVERT TO 8-BIT RGB
                          if (eightbit==true)       {
                                run("RGB Color");
                                                                          }
if (single==true)             {
  saveAs("tiff", resultsDir+nodotname+"_C2.tif");
   IJ.log("File saved: " +nodotname+"_C2.tif");
                                          }
  close();

if (ChannelNo==3)                          {
        selectWindow("C3-"+orig);
              // CONVERT TO 8-BIT RGB
                          if (eightbit==true)             {
                                run("RGB Color");
                                                                                }
if (single==true)             {
  saveAs("tiff", resultsDir+nodotname+"_C3.tif");
   IJ.log("File saved: " +nodotname+"_C3.tif");
                                          }
  close();
                                                                      }

  selectWindow(orig);

              // CONVERT TO 8-BIT RGB
                          if (eightbit==true)       {
                                run("RGB Color");
                                                                          }
if (merge==true)              {
  saveAs("tiff", resultsDir+nodotname+"_merge.tif");
   IJ.log("File saved: " +nodotname+"_merge.tif");
                                          }
run("Close All");
```

```
                                                                                                  }
                              } //END OF CZI LOOP

                                                                                                  }
//END OF FULL LOOP

getDateAndTime(year, month, dayOfWeek, dayOfMonth, hour, minute, second, msec);

twodigityear = year - 2000;

if (dayOfMonth<10) {
dayOfMonth = "0" + toString(dayOfMonth);
}

month = month+1;
if (month<10) {
month = "0" + month;
}

if (hour<10) {
hour = "0" + hour;
}

if (minute<10) {
minute = "0" + minute;
}

   ts = toString(dayOfMonth) + "-" + month + "-" + year + " " + hour + ":" + minute;
   IJ.log("FINISHED processing: " + ts);
selectWindow("Log");
saveAs("Text",
resultsDir+twodigityear+month+toString(dayOfMonth)+"_"+hour+"h"+minute+"m"+"_Log.txt");
showMessage( "Fin","FINISHED Processing");
```

### Macro 2 "GEN_Morphometrics.ijm"

```
var Parent_ROI = "1";

macro "Enter Parent ROI name [0]"	{
			roiManager("UseNames", "true");
			roiManager("select", 0);
			roiManager("Show All with labels");
	Dialog.create("Enter Parent ROI name");
	Dialog.addMessage("Enter Parent ROI name");
	Dialog.addString("Parent ROI:", Parent_ROI, 12);
	Dialog.show();
	Parent_ROI = Dialog.getString();
	setTool("freeline");
								} //Macro
macro "Line ROI trace [9]"	{
			roiManager("Add");
			nroi = roiManager("count");
			roiManager("select", nroi-1);
			roiManager("rename",Parent_ROI+"_line");
			roiManager("select", 0);
			roiManager("UseNames", "true");
			roiManager("Show All without labels");
						} //Macro

macro "Fibre Trace [7]"{
//"Fibre_Trace" macro for ImageJ
//Adds ROIs to ROI manager but doesn't allow overlaps. New ROIs are trimmed to the borders of existing ROIs.
roiManager("Add");
				//waitForUser( "Pause","Check selection");
newname = roiManager("count");
roiManager("select", newname-1);
roiManager("rename",newname);
roiManager("Show All with labels");
nroi = roiManager("count");
//print("nroi:");
//print(nroi);
if(nroi>1)	{
roiManager("Deselect");

//MAKE ARRAY OF ALL FIBRES
Fibres=newArray(0);
for (i=0;i<roiManager("count");i++)	{
roiManager("Select",i);
currentName=getInfo("roi.name");
if (matches(currentName,".*-.*"))	{
								//do nothing!
								}else

								{
	Fibres = Array.concat(Fibres,i);
								}

}
//print("Fibres");
//Array.print(Fibres);
```

```
//ARRAY OF ALL FIBRES EXCEPT THE LAST ONE
if(Fibres.length>2)        {
  FibresMinusOne = Array.trim(Fibres,Fibres.length-1);
            //print("FibresMinusOne");
            //Array.print(FibresMinusOne);

//GROUP ALL FIBRES EXCEPT THE LAST ONE
roiManager("select", FibresMinusOne);
//print("Fibres");
//Array.print(Fibres);
roiManager("Combine");
roiManager("Add");
roiManager("Deselect");
                                    } else    {roiManager("select", Fibres[0]);
                                    //waitForUser( "Pause","Check selection");
                                    roiManager("Add");
                                    roiManager("Deselect");

                                                            }

//SELECT GROUPED PREVIOUS ROIs and THE NEW ROI
if(Fibres.length>2)        {
array_last2 = newArray(nroi,nroi-1);
                                    } else    {array_last2 = newArray(1,2);}
//print("array_last2:");
//Array.print(array_last2);
roiManager("select", array_last2);
                                    //waitForUser( "Pause","Check selection");
roiManager("AND");
   type = selectionType();
            //DELETES COMBINED GROUP FROM ROI MANAGER WHEN NO OVERLAP
   if (type==-1) {
            roiManager("Deselect");
                        roiManager("select", nroi);
                        roiManager("Delete");
                        //waitForUser("Pause","No overlap");
allcount = roiManager("count");
roiManager("select", allcount-1);
roiManager("rename",Fibres.length);
            roiManager("Show All with labels");
                  //waitForUser("Pause","Check ROI names");
                                                            }
      //DELETES COMBINED GROUP AND "AND" GROUP FROM ROI MANAGER
   if (type!=-1)  {
roiManager("Add");

a = nroi-1;
b = nroi+1;

array_for_XOR = newArray(a,b);
roiManager("select", array_for_XOR);
roiManager("XOR");
roiManager("Add");
roiManager("Deselect");

array_for_del = newArray(a,b,nroi);
```

```
roiManager("select", array_for_del);
    roiManager("Delete");
    roiManager("Deselect");
allcount = roiManager("count");
roiManager("select", allcount-1);
roiManager("rename",Fibres.length);
            roiManager("Show All with labels");
                 //waitForUser("Pause","Check ROI names");
                                                  }
}
}//macro


macro "Myofibril Trace [8]"{
//"Fibre_Trace" macro for ImageJ
//Adds ROIs to ROI manager but doesn't allow overlaps. New ROIs are trimmed to the borders of existing ROIs.
roiManager("Add");
Roi.setFillColor("#33ff0000");
newname = roiManager("count");
roiManager("select", newname-1);
roiManager("rename",newname+"-1");
roiManager("Show All with labels");
nroi = roiManager("count");
//print("nroi:");
//print(nroi);
if(nroi>1)         {
roiManager("Deselect");

//MAKE ARRAY OF ALL OF THE INDICES IN THE ROI MANAGER
array1 = newArray("0");;
for (i=1;i<roiManager("count");i++){
       array1 = Array.concat(array1,i);
}
//print("array1");
//Array.print(array1);
//SELECT LAST TWO ROIs IN THE LIST
          //ARRAY OF LAST TWO VALUES
if(nroi>0)         {
  lasttwo = Array.reverse(array1);
      //Array.print(lasttwo);
      //waitForUser( "Pause","Check array");
  lasttwo = Array.trim(lasttwo,2);
      //Array.print(lasttwo);
      //waitForUser( "Pause","Check array");
  lasttwo = Array.reverse(lasttwo);
      //Array.print(lasttwo);
      // waitForUser( "Pause","Check array");
  roiManager("select", lasttwo);
      // waitForUser( "Pause","Check selection");

roiManager("And");
   type = selectionType();
   if (type!=-1)  {
roiManager("Add");
roiManager("Deselect");
roiManager("select", nroi-1);
```

```
roiManager("delete");
roiManager("select", nroi-1);
newname = roiManager("count");
roiManager("rename",newname+"-1");
        roiManager("Show All with labels");

//SHUFFLE LAST ROI TO THE BEGINNING
roicount = roiManager("count");
for (j=0;j<roicount-1;j++) {
            roiManager("Select",0);
            origLabel=getInfo("roi.name");
            roiManager("Add");//creates the same selection at bottom of list
            roiManager("Delete");//deletes the old selection
            roiManager("Select",roicount-1);//selects the new entry at bottom of list
            roiManager("Rename",origLabel);
      }
             //waitForUser("Pause","Check ROIs");

//GET NUMBER OF LAST FIBRE
            roiManager("Select",roicount-1);
            fibreNumber=getInfo("roi.name");

//COUNT NUMBER OF MYOFIBRILS ALREADY ASSOCIATED WITH FIBRE
fibrilcount = 1;
for (i=0;i<roiManager("count");i++)     {
                    roiManager("Select",i);
                    roiName=getInfo("roi.name");
                    if(startsWith(roiName,fibreNumber+"-")){fibrilcount = fibrilcount+1;}
                                                                        }
//RENAME FIBRIL ACCORDINGLY
            roiManager("Select",0);
            roiManager("Rename",fibreNumber+"-"+fibrilcount);
            Roi.setFillColor("#33ff0000");
                                        } else {
roiManager("Deselect");
roiManager("select", nroi-1);
//waitForUser("Pause","check selection for deletion");
roiManager("delete");
waitForUser("Pause","Select fibril inside last fibre!");
                                        }
}

}//macro
```

### Macro 3 “GEN_Split_to_green_and_blue_highlight_saturated.ijm”

```
// SPECIFYING DIRECTORY, LISTING INPUT FILES AND MAKING RESULTS FOLDER
Imagepath = getDirectory("Choose a Directory containing lsm/czi files");
//Imagepath = "/Users/thomas.hall/Desktop/test2/";
list = getFileList(Imagepath);

//SETTING UP LOG FILE
print("\\Clear");
getDateAndTime(year, month, dayOfWeek, dayOfMonth, hour, minute, second, msec);

if (dayOfMonth<10) {
dayOfMonth = "0" + toString(dayOfMonth);
}

month = month+1;
if (month<10) {
month = "0" + month;
}

if (hour<10) {
hour = "0" + hour;
}

if (minute<10) {
minute = "0" + minute;
}

   ts = toString(dayOfMonth) + "-" + month + "-" + year + " " + hour + ":" + minute;
   IJ.log("STARTED macro: Create merged for ROI selection.ijm, " + ts);

//SPECIFYING FILE EXTENSIONS
ImageEnd = ".lsm";
ImageEnd1 = ".czi";

//COUNT FILES TO PROCESS
  count = 0;
  for (i=0; i<list.length; i++) {
                if(endsWith(list[i],ImageEnd)||endsWith(list[i],ImageEnd1))       {
          count++;}

        }//count
         print("Total files in folder " + list.length);
                         print("Number of image files to process: " +count);


// DIALOGUE BOX FOR INPUT
                subfolder = "Split";
                Dialog.create("Specify Variables");
                        Dialog.addString("New subfolder:", subfolder, 30);
                        Dialog.addCheckbox("Enhance Contrast (0.35% saturated pixels)", true);
                        Dialog.addCheckbox("Highlight saturated pixels in red", true);
                        Dialog.addCheckbox("Run in batch mode", false);
                        Dialog.show();
                subfolder = Dialog.getString();
                EnhCont = Dialog.getCheckbox();
                Highlight = Dialog.getCheckbox();
```

```
                    Batch = Dialog.getCheckbox();

     resultsDir = Imagepath+subfolder+"/";
     File.makeDirectory(resultsDir);

    IJ.log("Input folder: " + Imagepath);
    IJ.log("Subfolder: " + subfolder);
                    if(EnhCont==1){
                    EnhCont_txt = "YES";
                    } else {EnhCont_txt = "NO";}
    IJ.log("Enhance Contrast (0.35% saturated pixels): " + EnhCont_txt);
                    if(Highlight==1)          {
                    Highlight_txt = "YES";
                    } else {Highlight_txt = "NO";}
    IJ.log("Highlight saturated pixels in red: " + Highlight_txt);

setBatchMode(Batch);

//LOOP

for (i=0; i<list.length; i++)                                        {

// OPEN FILE

 if (endsWith(list[i],ImageEnd)||endsWith(list[i],ImageEnd1))                              {
run("Bio-Formats Importer", "open=" +Imagepath+list[i]+" color_mode=Default view=Hyperstack
stack_order=XYCZT");
  orig = getTitle();
  IJ.log("File opened: " + orig + "         file " + (i+1) + " of " + count+1);
 if (endsWith(orig,ImageEnd))
  {nodotname = replace(orig,ImageEnd, "");};
 if (endsWith(orig,ImageEnd1))
  {nodotname = replace(orig,ImageEnd1, "");};
//    IJ.log("nodotname: " + nodotname);
//waitForUser("Check log");
//CLEAR ROI MANAGER
nroi = roiManager("count");
roiManager("Deselect");
for (j=0; j<nroi; j++){
roiManager("Select", 0);
roiManager("Delete");
}


run("Split Channels");
//waitForUser("Check log");
// SELECT CHANNEL 1 AND SET AS GREEN, CREATE ROI FROM SATURATED PIXELS
selectWindow("C1-"+orig);
channel1 = getTitle();
//run("Green");
val1=65535;
val2=65535;
setThreshold(val1,val2,"no color");
run("Create Selection");
//waitForUser("Check selection");
Type = selectionType();
if (Type!=-1)    {
```

```
roiManager("Add");
                                    }
//waitForUser("Check ROI manager");
run("8-bit");
                                    if (EnhCont==true)              {
                                             run("Select None");
                                             //waitForUser("before EnhCont");
                                             run("Enhance Contrast", "saturated=0.35");
                                             //waitForUser("after EnhCont");
                                                                                              }

// SELECT CHANNEL 2 AND SET AS BLUE, CREATE ROI FROM SATURATED PIXELS
selectWindow("C2-"+orig);
channel2 = getTitle();
//run("Blue");
val1=65535;
val2=65535;
setThreshold(val1,val2,"no color");
run("Create Selection");
Type = selectionType();
if (Type!=-1)    {
roiManager("Add");
                                    }
run("8-bit");
                                    if (EnhCont==true)              {
                                             run("Select None");
                                             //waitForUser("before EnhCont");
                                             run("Enhance Contrast", "saturated=0.35");
                                             //waitForUser("after EnhCont");
                                                                                              }

//CREATE MERGED RGB 8-Bit IMAGE
//  IJ.log("C2-"+orig);
//  IJ.log("C1-"+orig);
//waitForUser("Check log");
run("Merge Channels...", "green="+channel1+" blue="+channel2+" create keep ignore");
run("RGB Color");
//waitForUser("Merge");

//CONVERT SATURATED PIXELS TO RED USING ROIs
         //Select all ROIs in ROI manager
                                             if (Highlight==true)             {
                  allROIs = newArray("0");
                  for (k=1;k<roiManager("count");k++)   {
      allROIs = Array.concat(allROIs,k);
                                                                                                       }
                  if (roiManager("count")>1)       {
                  roiManager("select", allROIs);
                  roiManager("Combine");
                  roiManager("Add");
                                                                                    }
                  //select last ROI in ROI manager
                  if (roiManager("count")>0)       {
                  roiManager("Select", (roiManager("count")-1));
                  setForegroundColor(255, 0, 0);
                  roiManager("Fill");
                                                                                    }
```

```
                                                                                                    }
//waitForUser("Final Image");

//SAVE MERGED TIF IN SUBFOLDER WITH SAME NAME AS ORIGINAL
saveAs("tiff", resultsDir+orig);

//CLEAR ROI MANAGER
nroi = roiManager("count");
roiManager("Deselect");
for (j=0; j<nroi; j++){
roiManager("Select", 0);
roiManager("Delete");
}

//CLOSE ALL WINDOWS
    while (nImages>0) {
        selectImage(nImages);
        close();
  }
 }//END OF "OPEN" LOOP
}//END OF FOR LOOP
IJ.log("Finished!");
```

### Macro 4 “TT_GET_box_ROIs_for_amplitude_comparison.ijm”

```
// SPECIFYING DIRECTORY, LISTING INPUT FILES AND MAKING RESULTS FOLDER
Imagepath = getDirectory("Choose a Directory containing lsm/czi files");
//Imagepath = "/Users/thomas.hall/Desktop/test2/";
list = getFileList(Imagepath);

//SETTING UP LOG FILE
print("\\Clear");
getDateAndTime(year, month, dayOfWeek, dayOfMonth, hour, minute, second, msec);

if (dayOfMonth<10) {
dayOfMonth = "0" + toString(dayOfMonth);
}

month = month+1;
if (month<10) {
month = "0" + month;
}

if (hour<10) {
hour = "0" + hour;
}

if (minute<10) {
minute = "0" + minute;
}

   ts = toString(dayOfMonth) + "-" + month + "-" + year + " " + hour + ":" + minute;
   IJ.log("STARTED macro: Create merged for ROI selection.ijm, " + ts);

//SPECIFYING FILE EXTENSIONS
ImageEnd = ".lsm";
ImageEnd1 = ".czi";

//COUNT FILES TO PROCESS
  count = 0;
  for (i=0; i<list.length; i++) {
                if(endsWith(list[i],ImageEnd)||endsWith(list[i],ImageEnd1))       {
          count++;}

        }//count
         print("Total files in folder " + list.length);
                        print("Number of image files to process: " +count);


// DIALOGUE BOX FOR INPUT
                subfolder = "Split";
                Dialog.create("Specify Variables");
                        Dialog.addString("New subfolder:", subfolder, 30);
                        Dialog.addCheckbox("Enhance Contrast (0.35% saturated pixels)", true);
                        Dialog.addCheckbox("Highlight saturated pixels in red", true);
                        Dialog.addCheckbox("Run in batch mode", false);
                        Dialog.show();
                subfolder = Dialog.getString();
                EnhCont = Dialog.getCheckbox();
                Highlight = Dialog.getCheckbox();
```

```
                    Batch = Dialog.getCheckbox();

     resultsDir = Imagepath+subfolder+"/";
     File.makeDirectory(resultsDir);

    IJ.log("Input folder: " + Imagepath);
    IJ.log("Subfolder: " + subfolder);
                    if(EnhCont==1){
                    EnhCont_txt = "YES";
                    } else {EnhCont_txt = "NO";}
    IJ.log("Enhance Contrast (0.35% saturated pixels): " + EnhCont_txt);
                    if(Highlight==1)          {
                    Highlight_txt = "YES";
                    } else {Highlight_txt = "NO";}
    IJ.log("Highlight saturated pixels in red: " + Highlight_txt);

setBatchMode(Batch);

//LOOP

for (i=0; i<list.length; i++)                                          {

// OPEN FILE

 if (endsWith(list[i],ImageEnd)||endsWith(list[i],ImageEnd1))                            {
run("Bio-Formats  Importer",  "open="  +Imagepath+list[i]+"  color_mode=Default  view=Hyperstack
stack_order=XYCZT");
  orig = getTitle();
  IJ.log("File opened: " + orig + "        file " + (i+1) + " of " + count+1);
 if (endsWith(orig,ImageEnd))
  {nodotname = replace(orig,ImageEnd, "");};
 if (endsWith(orig,ImageEnd1))
  {nodotname = replace(orig,ImageEnd1, "");};
//    IJ.log("nodotname: " + nodotname);
//waitForUser("Check log");
//CLEAR ROI MANAGER
nroi = roiManager("count");
roiManager("Deselect");
for (j=0; j<nroi; j++){
roiManager("Select", 0);
roiManager("Delete");
}


run("Split Channels");
//waitForUser("Check log");
// SELECT CHANNEL 1 AND SET AS GREEN, CREATE ROI FROM SATURATED PIXELS
selectWindow("C1-"+orig);
channel1 = getTitle();
//run("Green");
val1=65535;
val2=65535;
setThreshold(val1,val2,"no color");
run("Create Selection");
//waitForUser("Check selection");
Type = selectionType();
if (Type!=-1)    {
```

```
roiManager("Add");
                                        }
//waitForUser("Check ROI manager");
run("8-bit");
                                        if (EnhCont==true)              {
                                                  run("Select None");
                                                  //waitForUser("before EnhCont");
                                                  run("Enhance Contrast", "saturated=0.35");
                                                  //waitForUser("after EnhCont");
                                                                                                  }

// SELECT CHANNEL 2 AND SET AS BLUE, CREATE ROI FROM SATURATED PIXELS
selectWindow("C2-"+orig);
channel2 = getTitle();
//run("Blue");
val1=65535;
val2=65535;
setThreshold(val1,val2,"no color");
run("Create Selection");
Type = selectionType();
if (Type!=-1)    {
roiManager("Add");
                                        }
run("8-bit");
                                        if (EnhCont==true)              {
                                                  run("Select None");
                                                  //waitForUser("before EnhCont");
                                                  run("Enhance Contrast", "saturated=0.35");
                                                  //waitForUser("after EnhCont");
                                                                                                  }

//CREATE MERGED RGB 8-Bit IMAGE
//  IJ.log("C2-"+orig);
//  IJ.log("C1-"+orig);
//waitForUser("Check log");
run("Merge Channels...", "green="+channel1+" blue="+channel2+" create keep ignore");
run("RGB Color");
//waitForUser("Merge");

//CONVERT SATURATED PIXELS TO RED USING ROIs
          //Select all ROIs in ROI manager
                                                  if (Highlight==true)              {
                    allROIs = newArray("0");
                    for (k=1;k<roiManager("count");k++)   {
      allROIs = Array.concat(allROIs,k);
                                                                                                            }
                    if (roiManager("count")>1)       {
                    roiManager("select", allROIs);
                    roiManager("Combine");
                    roiManager("Add");
                                                                                        }
                    //select last ROI in ROI manager
                    if (roiManager("count")>0)       {
                    roiManager("Select", (roiManager("count")-1));
                    setForegroundColor(255, 0, 0);
                    roiManager("Fill");
                                                                                        }
```

```
                                                                                    }
//waitForUser("Final Image");

//SAVE MERGED TIF IN SUBFOLDER WITH SAME NAME AS ORIGINAL
saveAs("tiff", resultsDir+orig);

//CLEAR ROI MANAGER
nroi = roiManager("count");
roiManager("Deselect");
for (j=0; j<nroi; j++){
roiManager("Select", 0);
roiManager("Delete");
}

//CLOSE ALL WINDOWS
      while (nImages>0) {
          selectImage(nImages);
          close();
  }
 }//END OF "OPEN" LOOP
}//END OF FOR LOOP
IJ.log("Finished!");
```

### Macro 5 “TT_perturbation_process.ijm”

```
//This macro takes line ROIS from expressing and control muscle fibres and compares the peaks (green caax channel) to look for peturbations.

//SETTING UP LOG FILE
print("\\Clear");
getDateAndTime(year, month, dayOfWeek, dayOfMonth, hour, minute, second, msec);

if (dayOfMonth<10) {
dayOfMonth = "0" + toString(dayOfMonth);
}

month = month+1;
if (month<10) {
month = "0" + month;
}

if (hour<10) {
hour = "0" + hour;
}

if (minute<10) {
minute = "0" + minute;
}

    ts = toString(dayOfMonth) + "-" + month + "-" + year + " " + hour + ":" + minute;
    IJ.log("STARTED macro: " + ts);

//setBatchMode(true);

//SET VARIABLES HERE FOR TWEAKING ALSO
oo = 0;
group = 6;//number of files for each group (gene)

// SPECIFYING DIRECTORY, LISTING INPUT FILES AND MAKING RESULTS FOLDER
        IJ.log("Choose a Directory containing input files");
    Imagepath = getDirectory("Choose a Directory containing image files");
    //Imagepath = "/Users/thomas.hall/Desktop/arse/";
        IJ.log("Choose a Directory containing ROI files");
    ROIpath = getDirectory("Choose a Directory containing ROI files");
        //ROIpath = "/Users/thomas.hall/Desktop/arse/ROIs/";
    list = getFileList(Imagepath);



// DIALOGUE BOX FOR INPUT
        expgroup = "TT";
        ImageEnd = ".czi";
        TTend = "_TT_box_ROI.zip";
        ControlEnd = "_control_TT_box_ROI.zip";
        LimitPeaks = true; //Used to calculate min and max for normalisation - limits averging to the fist x peaks/troughs
        PeakCalc = 6; //Number of peaks and trough to use
        subdir = "RawData";
        subdir1 = "SummaryData";
        minperiodum = 1.9; //actual sarcomere length is 1.9um
```

```
	rolling1 = "500";
	radius1 = "1";

	Dialog.create("Enter Parameters");
		Dialog.addString("Image Filenames end with:", ImageEnd, 30);
		Dialog.addString("TT ROI Filenames end with:", TTend, 30);
		Dialog.addString("Control ROI Filenames end with:", ControlEnd, 30);
		Dialog.addNumber("Period (sarcomere length in um)", minperiodum);
		Dialog.addCheckbox("Specify number of peaks/troughs for normalisation calculations",
false);
		Dialog.addNumber("Number of peaks/troughs to use:", PeakCalc);
		Dialog.addCheckbox("Use background subtraction", false);
		Dialog.addNumber("Rolling ball size in pixels", rolling1);
		Dialog.addCheckbox("Use median filter", true);
		Dialog.addNumber("Radius in pixels", radius1);
		Dialog.addCheckbox("Apply grayscale LUT", false);
	Dialog.show();

	ImageEnd = Dialog.getString();
	TTend = Dialog.getString();
	ControlEnd = Dialog.getString();
	minperiodum = Dialog.getNumber();
	LimitPeaks = Dialog.getCheckbox();
	if(LimitPeaks == 0) {LimitPeaksString = "NO";} else {LimitPeaksString = "YES";}
	PeakCalc = Dialog.getNumber();
	BGsub = Dialog.getCheckbox();
	if(BGsub == 0) {BGsubString = "NO";} else {BGsubString = "YES";}
	rolling1 = Dialog.getNumber();
	median = Dialog.getCheckbox();
	if(median == 0) {medianString = "NO";} else {medianString = "YES";}
	radius1 = Dialog.getNumber();
	gLUT = Dialog.getCheckbox();

//PRINT PARAMETERS TO LOG
IJ.log("USER DEFINED PARAMETERS:");
IJ.log("Minimum period in um: " + minperiodum);
IJ.log("Specify number of peaks/troughs for normalisation calculations: " + LimitPeaksString);
if(LimitPeaks==true)	{
IJ.log("Number of peaks/troughs to use for normalisation calculations: " + PeakCalc);
						};
IJ.log("Use background subtraction: " + BGsubString);
if(BGsub==true)		{
IJ.log("Rolling ball size in pixels: " + rolling1);
					};
IJ.log("Use median filter: " + medianString);
if(median==true)		{
IJ.log("Radius in pixels: " + radius1);
					};
//COUNT FILES TO PROCESS
  count = 0;
  for (i=0; i<list.length; i++) {
	   if (endsWith(list[i],ImageEnd)){
	   count++;
	}
     }
			print("Total files in folder " + list.length);
			print("Number of files to process: " +count);
```

```
//MAKING SUBDIRECTORIES FOR DATA SPIT
File.makeDirectory(Imagepath+"/"+subdir);
datapath = Imagepath+subdir+"/";
File.makeDirectory(Imagepath+"/"+subdir1);
summarypath = Imagepath+subdir1+"/";
File.makeDirectory(summarypath+"/"+"#Normalised_trace_data"+"/");
tracepath = summarypath+"/"+"#Normalised_trace_data"+"/";
File.makeDirectory(summarypath+"/"+"#Normalised_graphs"+"/");
graphpath = summarypath+"/"+"#Normalised_graphs"+"/";
File.makeDirectory(summarypath+"/"+"#Raw_graphs"+"/");
rawgraphpath = summarypath+"/"+"#Raw_graphs"+"/";
File.makeDirectory(summarypath+"/"+"#Raw_Peaks"+"/");
rawpeakspath = summarypath+"/"+"#Raw_Peaks"+"/";
File.makeDirectory(summarypath+"/"+"#Consolidated_Peaks"+"/");
conspeakspath = summarypath+"/"+"#Consolidated_Peaks"+"/";

// OPEN IMAGE FILE
    for (ww=0; ww<list.length; ww++) {
            if (endsWith(list[ww],ImageEnd)){
                            path = Imagepath+list[ww];//this way of opening supresses the dialogue box
for czi files
                            run("Bio-Formats Importer", "open=" +path+" color_mode=Default
view=Hyperstack stack_order=XYCZT use_virtual_stack");
                    orig = getTitle();
            //waitForUser( "Pause","Check open file");
            nodotname = replace(orig,ImageEnd, "");
                    IJ.log("Processing file: " + orig + " ("+(ww+1)+" of " + count + ")");

//GET SCALE
                    getPixelSize(unit, pixelWidth, pixelHeight);
                    onePixel_in_um = 1/pixelWidth;
                    one_um_in_Pixels = pixelWidth;
                    //print("Pixel size: " + "x= " + pixelWidth + " y= " + pixelHeight + " Unit= " + unit);
                    //print("1 pixel= " + onePixel_in_um + unit);
                    periodpx = round(minperiodum*onePixel_in_um);
                    //print("Min period in pixels = " + periodpx);

// TRANSFORMATIONS
        if (gLUT==true)         {
        run("Grays");
                                        }

        if (BGsub==true)        {
                run("Subtract Background...", "rolling=&rolling1 sliding");
                                        }

        if (median==true)       {
                run("Median...", "radius=&radius1");
                                                }


//SET UP LOOP HERE FOR THE CONTROL VS THE TT
    for (vv=0; vv<2; vv++)      {
if (vv == 0) {ROIend = ControlEnd; expgroup = "Control";} else //Control is processed first otherwise
can't normalise values to it
                            {ROIend = TTend; expgroup = "TT";}
```

```
                                        ROIfilename = nodotname + ROIend;

// CLEAR ROI MANAGER
     nroi = roiManager("count");
     roiManager("Deselect");

  for (j=0; j<nroi; j++){
    roiManager("Select", 0);
    roiManager("Delete");
  } // for j

// OPEN ROI FILE
 roiManager("Open", ROIpath + ROIfilename);
    roiManager("Select", 0);
     //waitForUser( "Pause","Check ROI");

//get coordinates of bottom of y line
    roiManager("Select", 1);
getSelectionCoordinates(xPoints,yPoints);
x = xPoints[0];
y = yPoints[0];
Array.reverse(xPoints);
Array.reverse(yPoints);
x1 = xPoints[0];
y1 = yPoints[0];
xoffset = x1-x;
yoffset = y1-y;

//print("xoffset = " + xoffset);
//print("yoffset = " + yoffset);

meanarray=newArray(0);

//Get coordinates of x line and set up loop for measurements in y
   roiManager("Select", 0);
            run("Interpolate", "interval=1"); // Edit>Selection>Interpolate
 getSelectionCoordinates(xPoints, yPoints);
//Round the points to the nearest pixel
            for (i=0; i<xPoints.length; i++) {
xRnd = round(xPoints[i]);
xPoints[i] = xRnd;
yRnd = round(yPoints[i]);
yPoints[i] = yRnd;
} //for i

            for (i=0; i<xPoints.length; i++) {
x = xPoints[i];
//print("xPoints = ");
//Array.print(xPoints);
y = yPoints[i];
//Array.print(yPoints);
Xend = x+xoffset;
Yend =  y+yoffset;
makeLine(x,y,Xend,Yend);
//waitForUser("check line selection for average");

yprofile = getProfile();
```

```
Array.getStatistics(yprofile, min, max, mean, stdDev);
//print("mean (" + (i+1) + ") = " + mean);
meanarray = Array.concat(meanarray, mean);
//print("Array of means: ");
//Array.print(meanarray);
        } // for i

//Set up series for Intensity Plot
n = meanarray.length;
xsequence = Array.getSequence(n);

//Find Maxima
                //Finds first peak within periodpx
firstPeak=Array.slice(meanarray, 0, periodpx);
//print("Array for finding first peak");
//Array.print(firstPeak);
rankPos_firstPeak = Array.rankPositions(firstPeak);//value of x for peak will be last value in this array
//print("Ranked positions");
//Array.print(rankPos_firstPeak);
firstx=rankPos_firstPeak[rankPos_firstPeak.length-1]; //-1 because arrays start with 0
//print("first x = " + firstx);
Array.sort(firstPeak);
//Array.print(firstPeak);//value of y for peak will be last value in this array
firsty=firstPeak[firstPeak.length-1]; //-1 because arrays start with 0
//print("first y = " + firsty);
//waitForUser("Check data for first maximum");
                //

        maxLocsY=newArray(0);
        maxLocsX=newArray(0);
Plot.create(expgroup +" Profile", "Distance in px", "Intensity");
Plot.setColor("red", "red");
Plot.add("line", xsequence, meanarray);
Plot.setFontSize(14);
Plot.show;
//waitForUser("Profile plotted");

//Find values of x and y for peaks
        done = false;
        for (jj= 0; jj < meanarray.length && !done; jj++){ //Finds max Y value within periodpx range
for x, using predicted next location of x
                if(jj==0) {predictx = firstx; lastx=firstx-periodpx;} else
                        {lastx = maxLocsX[maxLocsX.length-1];
                        predictx = lastx+periodpx;                                  }
                //print("predictx = " + predictx);
                //print("lastx = " + lastx);
                lower_window = predictx-(periodpx/2);
                if(lower_window <0) {lower_window = 0;}
                        //print("lower window = " + lower_window);
                upper_window = predictx+(periodpx/2);
                        //print("upper window = " + upper_window);
                nextPeak=Array.slice(meanarray, lower_window, upper_window);
                        //print("window array = ");
                //Array.print(nextPeak);
                rankPos_nextPeak = Array.rankPositions(nextPeak);//value of x for peak will be last
value in this array
                //Array.print(rankPos_nextPeak);
```

```
                    nextx=((rankPos_nextPeak[rankPos_nextPeak.length-1])+lower_window);      //sliced
array (+1) has to be added to last value of x
                    //print("next x = " + nextx);
                    Array.sort(nextPeak);
                    //Array.print(nextPeak);//value of y for peak will be last value in this array
                    nexty=nextPeak[nextPeak.length-1]; //-1 because arrays start with 0
                    //print("next y = " + nexty);
                              //waitForUser("check window array");

//Add values of x and Y to the peak arrays
                    maxLocsX = Array.concat(maxLocsX, nextx);
                    maxLocsY = Array.concat(maxLocsY, nexty);
toUnscaled(nextx, nexty); //don't know why you need this but it doesn't plot properly otherwise
makeOval(nextx-4, nexty-4, 8, 8); //creates circles at peaks, the minus makes it the middle of the circle
changeValues(0x0ffffff, 0x0ffffff, 0x33F33); // Makes circles light green
if(((maxLocsX[maxLocsX.length-1])+periodpx)>meanarray.length) {done = true;} //breaks loop;
          wait(100);
                    //waitForUser("Max data point plotted");
                                                                                                            }//for jj

//Find Minima
          minLocsX=newArray(0);
          minLocsY=newArray(0);

          done = false;
          for (jj= 0; jj < meanarray.length && !done; jj++){ //Finds min Y values between peaks
                    peak1 = maxLocsX[jj];
                    peak2 = maxLocsX[jj+1];
                    nextTrough=Array.slice(meanarray, peak1, peak2);
                    Array.getStatistics(nextTrough, min, max, mean, stdDev);
                    rankPos_nextTrough = Array.rankPositions(nextTrough);//value of x for peak will be
first value in this array
                    xmin = (rankPos_nextTrough[0]) + peak1;
                    Array.sort(nextTrough);
                    //Array.print(nextTrough);//value of y for peak will be first value in this array
                    ymin = nextTrough[0];

//Add values of x and Y to the trough arrays
                    minLocsX = Array.concat(minLocsX, xmin);
                    minLocsY = Array.concat(minLocsY, ymin);
toUnscaled(xmin, ymin); //don't know why you need this but it doesn't plot properly otherwise
makeOval(xmin-4, ymin-4, 8, 8); //creates circles at peaks, the minus makes it the middle of the circle
changeValues(0x0ffffff, 0x0ffffff, 0x000FF); // Makes circles dark blue
if((maxLocsX.length-1)<=(minLocsX.length)) {done = true;} //breaks loop;
          wait(100);
                    //waitForUser("Min data point plotted");
                              }//for jj

Array.show("Profile",xsequence,meanarray);//SAVE ALL UNPROCESSED DATA AND CLOSE WINDOWS
          selectWindow("Profile");
          saveAs("Results", datapath+nodotname+"_"+expgroup+"_profile.csv");
          selectWindow(nodotname+"_"+expgroup+"_profile.csv");
run("Close");
Array.show("Maxima",maxLocsX,maxLocsY);
          selectWindow("Maxima");
          saveAs("Results", datapath+nodotname+"_"+expgroup+"_maxima.csv");
          selectWindow(nodotname+"_"+expgroup+"_maxima.csv");
```

```
run("Close");
Array.show("Minima",minLocsX,minLocsY);
        selectWindow("Minima");
        saveAs("Results", datapath+nodotname+"_"+expgroup+"_minima.csv");
        selectWindow(nodotname+"_"+expgroup+"_minima.csv");
run("Close");
if(vv==0)       {
        control_raw_maxima = Array.copy(maxLocsY);
        control_raw_minima = Array.copy(minLocsY);
                        };
if(vv==1)       {
        TT_raw_maxima = Array.copy(maxLocsY);
        TT_raw_minima = Array.copy(minLocsY);
                        };
selectWindow(expgroup +" Profile");
saveAs("PNG", rawgraphpath+nodotname+"_"+expgroup+"_plot.png");
run("Close");

// CLEAR ROI MANAGER
     nroi = roiManager("count");
     roiManager("Deselect");

  for (j=0; j<nroi; j++){
    roiManager("Select", 0);
    roiManager("Delete");
  } // for j

//CALCULATE NORMALISED VALUES FROM CONTROL MINIMA AND MAXIMA

//CALCULATE ADJUSTED X VALUES
FirstPeakMinus = maxLocsX[0];
        maxLocsXminus = Array.copy(maxLocsX);
  for (j=0; j<maxLocsXminus.length; j++)        {
        maxLocsXminus[j] = maxLocsXminus[j] - (FirstPeakMinus-1);

        }//for j
Array.show("Maxima",maxLocsXminus,maxLocsX,maxLocsY);
//waitForUser("Check");

        minLocsXminus = Array.copy(minLocsX);
  for (j=0; j<minLocsXminus.length; j++)        {
        minLocsXminus[j] = minLocsXminus[j] - (FirstPeakMinus-1);

        }//for j
Array.show("Minima",minLocsXminus,minLocsX,minLocsY);
//waitForUser("Check");
        meanarrayminus = Array.slice(meanarray,FirstPeakMinus,meanarray[meanarray.length-1]);
                n = meanarrayminus.length;
                xsequenceminus = Array.getSequence(n+1);
                xsequenceminus = Array.slice(xsequenceminus,1,xsequenceminus.length);
Array.show("Profile",xsequence,meanarray,xsequenceminus,meanarrayminus);
//waitForUser("Check");

//CALCULATE ADJUSTED Y VALUES
if(expgroup == "Control") {
Array.getStatistics(maxLocsY, min, max, mean, stdDev);
YmaxMean = mean;
```

```
//print("YmaxMean = " + mean);
Array.getStatistics(minLocsY, min, max, mean, stdDev);
YminMean = mean;
//print("YminMean = " + mean);
if(LimitPeaks == true)   {//Limits the number of peaks/troughs for averaging to a specified number
defined by "PeakCalc"
        //print("PeakCalc= " + PeakCalc);
        maxLocsYlimited = Array.copy(maxLocsY);
        maxLocsYlimited = Array.slice(maxLocsYlimited, 0, PeakCalc);
        //print("YmaxMean (limited) array: ");
        //Array.print(maxLocsYlimited);
        Array.getStatistics(maxLocsYlimited, min, max, mean, stdDev);
        YmaxMean = mean;
        //print("YmaxMean (limited)= " + mean);

        minLocsYlimited = Array.copy(minLocsY);
        minLocsYlimited =Array.slice(minLocsYlimited, 0, PeakCalc);
        //print("YminMean (limited) array: ");
        //Array.print(minLocsYlimited);
        Array.getStatistics(minLocsYlimited, min, max, mean, stdDev);
        YminMean = mean;
        //print("YminMean (limited)= " + mean);
                                                        };//if LimitPeaks
//waitForUser("check data");
                                                        };//if expgroup

//SUBTRACT MEAN OF MINIMA TO NORMALISE TO ZERO
        meanarrayminuszeroed = Array.copy(meanarrayminus);
        maxLocsYzeroed = Array.copy(maxLocsY);
        minLocsYzeroed = Array.copy(minLocsY);
  for (j=0; j<meanarrayminuszeroed.length; j++) {
        meanarrayminuszeroed[j] = meanarrayminuszeroed[j] - (YminMean);

        }//for j
  for (j=0; j<maxLocsYzeroed.length; j++)        {
        maxLocsYzeroed[j] = maxLocsYzeroed[j] - (YminMean);

        }//for j
  for (j=0; j<minLocsYzeroed.length; j++)        {
        minLocsYzeroed[j] = minLocsYzeroed[j] - (YminMean);

        }//for j
//print("Zeroed");
//Array.print(maxLocsYzeroed);
//Array.print(minLocsYzeroed);

//CONVERT TO PERCENTAGE USING MEAN OF MAXIMA AS 100%
        meanarrayminuspercent = Array.copy(meanarrayminuszeroed);
        maxLocsYpercent = Array.copy(maxLocsYzeroed);
        minLocsYpercent = Array.copy(minLocsYzeroed);
  for (j=0; j<meanarrayminuspercent.length; j++) {
        meanarrayminuspercent[j] = (meanarrayminuspercent[j] / (YmaxMean-YminMean))*100;

        }//for j
  for (j=0; j<maxLocsYpercent.length; j++)       {
        maxLocsYpercent[j] = (maxLocsYpercent[j] / (YmaxMean-YminMean))*100;
```

```
	}//for j
  for (j=0; j<minLocsYpercent.length; j++)	{
	minLocsYpercent[j] = (minLocsYpercent[j] / (YmaxMean-YminMean))*100;

	}//for j
//print("Percentages");
//Array.print(maxLocsYpercent);
//Array.print(minLocsYpercent);

if (vv == 0)	{//vv is whether or not it is the control or TT
Plot.create("Normalised Plot", "Distance in px", "Intensity");
Plot.setColor("red", "red");
	x_control_line = Array.copy(xsequenceminus);
	y_control_line = Array.copy(meanarrayminuspercent);
	x_control_maxima = Array.copy(maxLocsXminus);
	y_control_maxima = Array.copy(maxLocsYpercent);
	x_control_minima = Array.copy(minLocsXminus);
	y_control_minima = Array.copy(minLocsYpercent);
//Array.print(y_control_maxima);
//Array.print(y_control_minima);
Plot.add("line", x_control_line, y_control_line);
Plot.setFontSize(14);
Plot.add("circles", x_control_maxima, y_control_maxima);
Plot.add("circles", x_control_minima, y_control_minima);
//Plot.show;
	//wait(500);
//waitForUser("check normalised control plot");
//selectWindow("Normalised Plot");
run("Close");
					} else {
Plot.create("Normalised Plot", "Distance in px", "Intensity");
Plot.setColor("red", "red");
Plot.add("line", x_control_line, y_control_line);
Plot.setFontSize(14);
Plot.add("circles", x_control_maxima, y_control_maxima);
Plot.add("circles", x_control_minima, y_control_minima);
//waitForUser("check normalised control plot");
Plot.setColor("blue", "blue");
Plot.add("line", xsequenceminus, meanarrayminuspercent);
Plot.setFontSize(14);
Plot.add("circles", maxLocsXminus, maxLocsYpercent);
Plot.add("circles", minLocsXminus, minLocsYpercent);
Plot.show;
	wait(500);
//waitForUser("check normalised control plot");
saveAs("PNG", graphpath+nodotname+"_"+"norm_plot.png");
run("Close");
y_controls =  Array.copy(y_control_line);
y_TTs = Array.copy(meanarrayminuspercent);
x_control_in_um =  Array.copy(x_control_line);
	 for (j=0; j<x_control_in_um.length; j++) {
	x_control_in_um[j] = (x_control_in_um[j]/onePixel_in_um);

	}//for j
x_TT_in_um =  Array.copy(xsequenceminus);
	 for (j=0; j<x_TT_in_um.length; j++) {
```

```
          x_TT_in_um[j] = (x_TT_in_um[j]/onePixel_in_um);

          }//for j
Array.show("Normalised Profiles",y_controls,y_TTs);
          selectWindow("Normalised Profiles");
          saveAs("Results", tracepath+nodotname+"_"+expgroup+".csv");
          selectWindow(nodotname+"_"+expgroup+".csv");
//waitForUser("pause");
run("Close");
//waitForUser("check normalised plot");
                                                  }//ELSE

//END OF CALCULATE NORMALISED VALUES SECTION

if (vv == 1)      {

//SAVES MAXIMA AND MINIMA (RAW VALUES) FOR TTs AND CONTROLS
Array.show("Peaks",control_raw_maxima,control_raw_minima,TT_raw_maxima,TT_raw_minima);
          selectWindow("Peaks");
          saveAs("Results", rawpeakspath+nodotname+"_Peaks.csv");
          selectWindow(nodotname+"_Peaks.csv");
//waitForUser("check data");
run("Close");

//SUMMARISES RESULTS FROM EACH BATCH OF SIX INTO ONE FILE (NASTY CODE)
oo = oo+1;//Loop for counting number of replicates
if(oo == 1) {     control_raw_maxima1 = Array.copy(control_raw_maxima);
                        control_raw_minima1 = Array.copy(control_raw_minima);
                        TT_raw_minima1 = Array.copy(TT_raw_minima);
                        TT_raw_maxima1 = Array.copy(TT_raw_maxima);}
if(oo == 2) {     control_raw_maxima2 = Array.copy(control_raw_maxima);
                        control_raw_minima2 = Array.copy(control_raw_minima);
                        TT_raw_minima2 = Array.copy(TT_raw_minima);
                        TT_raw_maxima2 = Array.copy(TT_raw_maxima);}
if(oo == 3) {     control_raw_maxima3 = Array.copy(control_raw_maxima);
                        control_raw_minima3 = Array.copy(control_raw_minima);
                        TT_raw_minima3 = Array.copy(TT_raw_minima);
                        TT_raw_maxima3 = Array.copy(TT_raw_maxima);}
if(oo == 4) {     control_raw_maxima4 = Array.copy(control_raw_maxima);
                        control_raw_minima4 = Array.copy(control_raw_minima);
                        TT_raw_minima4 = Array.copy(TT_raw_minima);
                        TT_raw_maxima4 = Array.copy(TT_raw_maxima);}
if(oo == 5) {     control_raw_maxima5 = Array.copy(control_raw_maxima);
                        control_raw_minima5 = Array.copy(control_raw_minima);
                        TT_raw_minima5 = Array.copy(TT_raw_minima);
                        TT_raw_maxima5 = Array.copy(TT_raw_maxima);}
if(oo == 6) {     control_raw_maxima6 = Array.copy(control_raw_maxima);
                        control_raw_minima6 = Array.copy(control_raw_minima);
                        TT_raw_minima6 = Array.copy(TT_raw_minima);
                        TT_raw_maxima6 = Array.copy(TT_raw_maxima);
Array.show("Consolidated
Peaks",control_raw_maxima1,control_raw_minima1,TT_raw_maxima1,TT_raw_minima1,control_raw_
maxima2,control_raw_minima2,TT_raw_maxima2,TT_raw_minima2,control_raw_maxima3,control_ra
w_minima3,TT_raw_maxima3,TT_raw_minima3,control_raw_maxima4,control_raw_minima4,TT_raw
_maxima4,TT_raw_minima4,control_raw_maxima5,control_raw_minima5,TT_raw_maxima5,TT_raw_
minima5,control_raw_maxima6,control_raw_minima6,TT_raw_maxima6,TT_raw_minima6);
          selectWindow("Consolidated Peaks");
```

```
            saveAs("Results", conspeakspath+nodotname+"_Consolidated_Peaks.csv");
            IJ.log("File saved: " +nodotname+"_Consolidated_Peaks.csv");
            //waitForUser("check data");
run("Close");
oo = 0;
                                }


//CLOSES IMAGE FILE
selectWindow(nodotname+ImageEnd);
run("Close All");
IJ.log("File closed: " + orig);
                                        }
                                                                        }//END OF vv LOOP (GOES TWICE -
FOR EACH ROI, THEN CLOSES FILE)

}// if endswith loop
}//for ww (open file loop)

//FINISHING LOG FILE AND EXITING MACRO
getDateAndTime(year, month, dayOfWeek, dayOfMonth, hour, minute, second, msec);
twodigityear = year - 2000;

if (dayOfMonth<10)      {
dayOfMonth = "0" + toString(dayOfMonth);
                                                }

month = month+1;
if (month<10)                   {
month = "0" + month;
                                                }

if (hour<10)                    {
hour = "0" + hour;
                                                }

if (minute<10)          {
minute = "0" + minute;
                                                }

   ts = toString(dayOfMonth) + "-" + month + "-" + year + " " + hour + ":" + minute;
   IJ.log("FINISHED macro: " + ts);

selectWindow("Log");
saveAs("Text",
datapath+twodigityear+month+toString(dayOfMonth)+"_"+hour+"h"+minute+"m"+"_Log.txt");
run("Close");//have to use run here for some reason to close a non-image window
showMessage( "Fin","FINISHED Processing");

exit();
```

**Macro 6 "TT_GET_circle_ROIs_for_localisation_comparison.ijm"**

```
//Tom Hall 26th July 2016

//FIRST install the "Put circle" macro for making circle ROIs from a point selection. Also specify the circle diameter here.
                    waitForUser( "Pause","Install 'Put circle' macro");

// SPECIFYING DIRECTORY, LISTING INPUT FILES AND MAKING RESULTS FOLDER
          Imagepath = getDirectory("Choose a Directory containing input files");
          list = getFileList(Imagepath);

//NEED TO SPECIFY AN INITIAL STRING FOR nodotname WHICH WILL GET REPLACED
nodotname = "xxxxx";

// FIXED VARIABLES
          TTend = "_TTs_ROI.zip";
          SarcolemmaEnd = "_Sarcolemma_ROI.zip";
          BetweenEnd = "_between_TTs_ROI.zip";

// DIALOGUE BOX FOR INPUT
                    filecounter = 0;
                    subfolder = "#localisation_ROIs";
                    ImageEnd = ".tif";
                    ROIend = "_ROI.zip";
                    pxdiameter=0.25;
                    zoom=200;
                    Dialog.create("Specify Variables");
                              Dialog.addString("New subfolder:", subfolder, 30);
                              Dialog.addString("File extension of input files:", ImageEnd, 30);
                              Dialog.addString("File extension for ROI files:", ROIend, 30);
                              Dialog.addCheckbox("Edit Existing ROI files?", false);
                              Dialog.addCheckbox("Open unprocessed only?", true);
                              Dialog.addNumber("Circle diameter (um):", pxdiameter);
                              Dialog.addNumber("Apply zoom? (%):", zoom);
                              Dialog.show();
                    subfolder = Dialog.getString();
                    ImageEnd = Dialog.getString();
                    ROIend = Dialog.getString();
                    EditROI = Dialog.getCheckbox();
                    Unprocessed = Dialog.getCheckbox();
                    pxdiameter = Dialog.getNumber();
                    zoom = Dialog.getNumber();

                    roiDir = Imagepath+subfolder+"/";
                    File.makeDirectory(roiDir);

//COUNT FILES TO PROCESS
  count = 0;
  for (i=0; i<list.length; i++) {
            if (endsWith(list[i],ImageEnd)){
            count++;
        }
    }
                              print("Total files in folder " + list.length);
                              print("Number of image files to process: " +count);
                              Array.print(list);
```

```
// OPEN IMAGE FILE
     for (i=0; i<list.length; i++) {
          print("Current file: " + list[i]);
               if (endsWith(list[i],ImageEnd))   {
                    orig = list[i];
                    nodotname = replace(orig,ImageEnd, "");
                                                                  }
            if
((endsWith(list[i],ImageEnd)&&(!File.exists(roiDir+nodotname+"_Sarcolemma"+ROIend)))||(endsWith(list[i],ImageEnd)&&(File.exists(roiDir+nodotname+"_Sarcolemma"+ROIend))&&Unprocessed==false)){
                    path = Imagepath+list[i];//this way of opening supresses the dialogue box for czi files
                              run("Bio-Formats Importer", "open=" +path+" color_mode=Default view=Hyperstack stack_order=XYCZT");
            orig = getTitle();
            nodotname = replace(orig,ImageEnd, "");
            ROIfilename = nodotname + ROIend;
            filecounter = filecounter+1;

               getPixelSize(unit, pixelWidth, pixelHeight);
               pxradius = pxdiameter/2;
               umdiameter = pxdiameter/pixelWidth;
               umradius = umdiameter/2;

//APPLY ZOOM
getDimensions(width, height, channels, slices, frames);
run("Set... ", "zoom=" +zoom);

// CLEAR ROI MANAGER
     nroi = roiManager("count");
     roiManager("Deselect");

  for (j=0; j<nroi; j++){
    roiManager("Select", 0);
    roiManager("Delete");
  } // for j

//OPEN EXISTING TT ROI FILE
//SELECT TTs
if ((EditROI==true)&&(File.exists(roiDir+nodotname+"_TTs"+ROIend))) {
          roiManager("Open", roiDir+nodotname+"_TTs"+ROIend);
                    //make overlay circles from existing ROIs
                 nroi = roiManager("count");
                 roiManager("Deselect");
                         for (j=0; j<nroi; j++)    {
                    roiManager("Select", j);
                         setForegroundColor(255, 255, 255);
                         run("Fill", "slice");
                                                                  }

                                   }//EditROI
               Stack.setChannel(1);
               waitForUser(i + " of " + count,"Select TT point ROIs using '6'");
                    roiManager("deselect");
//SAVE ROI LIST (ONLY IF THERE IS A SELECTION)
                    nroi = roiManager("count");
```

```
                    if(nroi>0)      {
                    roiManager("save", roiDir+nodotname+"_TTs"+ROIend);
                                }
            // CLEAR ROI MANAGER
                    nroi = roiManager("count");
                    roiManager("Deselect");

                    for (j=0; j<nroi; j++)              {
                    roiManager("Select", 0);
            //ADD EACH TO OVERLAY
                    run("Add Selection...");
                    roiManager("Delete");
                                                            } // for j


            //SELECT POINTS IN BETWEEN TTs
            if      ((EditROI==true)&&(File.exists(roiDir+nodotname+"_between_TTs"+ROIend)))
        {
        roiManager("Open", roiDir+nodotname+"_between_TTs"+ROIend);
                    //make overlay circles from existing ROIs
                nroi = roiManager("count");
                roiManager("Deselect");
                        for (j=0; j<nroi; j++)  {
                    roiManager("Select", j);
                        setForegroundColor(255, 255, 255);
                        run("Fill", "slice");
                                                    }

                                }//EditROI
            Stack.setChannel(2);
            waitForUser(i+1 + " of " + count,"Select In between point ROIs using '6'");
                    roiManager("deselect");
//SAVE ROI LIST (ONLY IF THERE IS A SELECTION)
                    nroi = roiManager("count");
                    if(nroi>0)      {
                    roiManager("save", roiDir+nodotname+"_between_TTs"+ROIend);
                                }
            // CLEAR ROI MANAGER
                        nroi = roiManager("count");
                    roiManager("Deselect");

                    for (j=0; j<nroi; j++)              {
                    roiManager("Select", 0);
            //ADD EACH TO OVERLAY
                    run("Add Selection...");
                    roiManager("Delete");
                                                            } // for j

            //SELECT SARCOLEMMA

                        if
((EditROI==true)&&(File.exists(roiDir+nodotname+"_Sarcolemma"+ROIend)))  {
        roiManager("Open", roiDir+nodotname+"_Sarcolemma"+ROIend);
                    //make overlay circles from existing ROIs
                nroi = roiManager("count");
                roiManager("Deselect");
                        for (j=0; j<nroi; j++)  {
```

```
                    roiManager("Select", j);
                            setForegroundColor(255, 255, 255);
                            run("Fill", "slice");
                                                                                }

                                            }//EditROI
            Stack.setChannel(3);
            waitForUser(i+1 + " of " + count,"Select Sarcolemma point ROIs using '6'");
                    roiManager("deselect");


//SAVE ROI LIST (ONLY IF THERE IS A SELECTION)
                    nroi = roiManager("count");
                    if(nroi>0)          {
                    roiManager("save", roiDir+nodotname+"_Sarcolemma"+ROIend);
                                            }
            // CLEAR ROI MANAGER
                    nroi = roiManager("count");
                    roiManager("Deselect");

                    for (j=0; j<nroi; j++)                      {
                    roiManager("Select", 0);
            //ADD EACH TO OVERLAY
                    run("Add Selection...");
                    roiManager("Delete");
                                                                                        } // for j

            close();

 } //if
}// for i
```

**Macro 7 "GEN_Put_circle.ijm"**

```
macro "Put circle [6]"{
	if (!startsWith(getInfo("command.name"), "^"))
		exit("Use keyboard shortcut '6'");
				if (selectionType!=10)		{
			showMessage("Make point selection");
								}
		if (selectionType==10)			{
			pxdiameter=0.25;//Specify the diameter of the circle in um here
			getPixelSize(unit, pixelWidth, pixelHeight);
			umdiameter = pxdiameter/pixelWidth;
			getSelectionCoordinates(xpoint, ypoint);
			//IJ.log("x: " + xpoint[0] + "    y: " + ypoint[0]);
			makeOval(xpoint[0]-umdiameter,    ypoint[0]-umdiameter,    umdiameter*2,
umdiameter*2);
			setForegroundColor(255, 255, 255);
			run("Fill", "slice");
			roiManager("Add");
			roiManager("Show All with labels");
			roiManager("Show All");
						}

}//macro
```

### Macro 8 “TT_localisation_comparison_process.ijm”

```
print("\\Clear");
// SPECIFYING DIRECTORY, LISTING INPUT FILES AND MAKING RESULTS FOLDER
    Imagepath = getDirectory("Choose a Directory containing image files");
    //Imagepath = "/Users/thomas.hall/Desktop/test/";
    ROIpath = getDirectory("Choose a Directory containing ROI files");
        //ROIpath = "/Users/thomas.hall/Desktop/test/ROIs/";
    list = getFileList(Imagepath);

//NEED TO SPECIFY AN INITIAL STRING FOR nodotname WHICH WILL GET REPLACED
nodotname = "xxxxx";

//SET VARIABLES HERE
//Sarcolemma vs TT
        row = 0;
        MaxIntensityArray = newArray(0);
        ImageEnd = ".czi";
        Channels = 2;
        TTend = "_TTs_ROI.zip";
        SarcolemmaEnd = "_Sarcolemma_ROI.zip";
        BetweenEnd = "_between_TTs_ROI.zip";
        FilterRad = 1;

        Dialog.create("Specify Variables");
                                    //Channel = getDirectory("Choose a Directory containing ROI
files");
                Dialog.addString("Image Filenames end with:", ImageEnd, 30);
                Dialog.addString("TT ROI Filenames end with:", TTend, 30);
                Dialog.addString("Sarcolemma ROI Filenames end with:", SarcolemmaEnd, 30);
                Dialog.addString("Between TT ROI Filenames end with:", BetweenEnd, 30);
                Dialog.addCheckbox("Use median filter", true);
                Dialog.addNumber("Radius for filter:", FilterRad);
                Dialog.addCheckbox("Run in batch mode", false);
                Dialog.show();
        ImageEnd = Dialog.getString();
        TTend = Dialog.getString();
        SarcolemmaEnd = Dialog.getString();
        Median = Dialog.getCheckbox();
        FilterRad = Dialog.getNumber();
        Batch = Dialog.getCheckbox();

setBatchMode(Batch);

//COUNT FILES TO PROCESS
  count = 0;
  for (i=0; i<list.length; i++) {
          if (endsWith(list[i],ImageEnd)){
          count++;
        }
    }
                        print("Total files in folder " + list.length);
                        print("Number of image files to process: " +count);


// OPEN IMAGE FILE
    for (ww=0; ww<list.length; ww++) {
```

```
                    if (endsWith(list[ww],ImageEnd))          {
                            orig = list[ww];
                            nodotname = replace(orig,ImageEnd, "");
                                                                                        }

            if
((endsWith(list[ww],ImageEnd))&&(File.exists(ROIpath+nodotname+"_Sarcolemma_ROI.zip"))){
                            path = Imagepath+list[ww];//this way of opening supresses the dialogue box
for czi files
                            run("Bio-Formats    Importer",    "open="    +path+"    color_mode=Default
view=Hyperstack stack_order=XYCZT");
                    orig = getTitle();
                    nodotname = replace(orig,ImageEnd, "");
                                    IJ.log("Processing file: " + orig + " ("+(ww+1)+" of " + count + ")");
if(Median==true)          {
run("Median...", "radius=FilterRad stack");
                                              }
//LOOP FOR THE SARCOLEMMA VS THE TT
   for (m=0; m<Channels; m++) {
   for (vv=0; vv<3; vv++)       {
if (vv == 0)      {ROIend = SarcolemmaEnd; expgroup = "Sarcolemma";}
if (vv == 1)      {ROIend = TTend; expgroup = "TT";}
if (vv == 2)      {ROIend = BetweenEnd; expgroup = "Between_TTs";}
                                    ROIfilename = nodotname + ROIend;

// CLEAR ROI MANAGER
   nroi = roiManager("count");
   roiManager("Deselect");

  for (j=0; j<nroi; j++){
    roiManager("Select", 0);
    roiManager("Delete");
  } // for j

// OPEN ROI FILE
roiManager("Open", ROIpath + ROIfilename);
//IJ.log("Processing ROI: " + ROIfilename);
MaxIntensityArray = newArray(0);
  //GET ALL INTENSITY VALUES FROM ROIs INTO AN ARRAY AND CALCULATE THE Max
   nroi = roiManager("count");
   roiManager("Deselect");
  for (j=0; j<nroi; j++){
    roiManager("Select", j);
                    Stack.setChannel(m+1);
                                              //waitForUser( "Pause","Check channel selection");
                                            List.setMeasurements;
                                            MaxIntensity = List.getValue("Max");

          //print("MaxIntensity:");
                                            //print(MaxIntensity);
                                            MaxIntensityArray                                        =
Array.concat(MaxIntensityArray,MaxIntensity);
                                                      } // for j
//print("MaxIntensityArray:");
//Array.print(MaxIntensityArray);
Array.getStatistics(MaxIntensityArray, min, max, mean, stdDev);
                                            //IJ.log("Mean of Max Intensities: " + mean);
```

```
                                        //waitForUser( "Pause","Check log window");
setResult("Image_file", row, orig);
    roiManager("Deselect");
    run("Select All");
                List.setMeasurements;
                MaxImageIntensity = List.getValue("Max");
                MinImageIntensity = List.getValue("Min");
                MedianImageIntensity                    =                   ((MaxImageIntensity-
MinImageIntensity)/2)+MinImageIntensity;
setResult("Ch_"+(m+1)+"_Max_Image_Intensity", row, MaxImageIntensity);
setResult("Ch_"+(m+1)+"_Min_Image_Intensity", row, MinImageIntensity);
setResult("Ch_"+(m+1)+"_Median_Image_Intensity", row, MedianImageIntensity);
setResult("Ch_"+(m+1)+"_"+expgroup, row, mean);
     }// for m (Channels)
// CLEAR ROI MANAGER
    nroi = roiManager("count");
    roiManager("Deselect");

  for (j=0; j<nroi; j++){
   roiManager("Select", 0);
   roiManager("Delete");
                                                } // for j

    }//for vv (ROI manager open)
row = row+1;

  //CLOSES IMAGE FILE
selectWindow(nodotname+ImageEnd);
run("Close All");
IJ.log("File closed: " + orig);
                                }// if endswith ww
    }//for ww
```

**Python script “ROI to .svg”**

```
import sys
from java.awt import Color
from java.io import File
from ij import IJ
from ij.gui import ShapeRoi
from ij.plugin.frame import RoiManager
from org.jfree.graphics2d.svg import SVGGraphics2D, SVGUtils

imp = IJ.getImage()
rm = RoiManager.getInstance()
if rm is None:
	print 'None'
	sys.exit()

# convert ROIs in RoiManager to an array of shapeRois
jrois = rm.getRoisAsArray()
srois = [ShapeRoi(jroi) for jroi in jrois]

# http://www.jfree.org/jfreesvg/javadoc/
g2 = SVGGraphics2D(imp.getWidth(), imp.getHeight())
g2.setPaint(Color.BLACK)
px = 0.0
py = 0.0
for sroi in srois:
	g2.translate(px*-1, py*-1)
	px = sroi.getBounds().x
	py = sroi.getBounds().y
	g2.translate(px, py)
	g2.draw(sroi.getShape())

se = g2.getSVGElement()

# writing the file
path = ">>INSERT PATH FOR EXPORT HERE <<"
SVGUtils.writeToSVG(File(path), se)
```
